# Intrinsic diving reflex induces potent antioxidative response by activation of NRF2 signaling

**DOI:** 10.1101/2024.02.12.579910

**Authors:** Keren Powell, Steven Wadolowski, Willians Tambo, Joshua J. Strohl, Daniel Kim, Justin Turpin, Yousef Al-Abed, Michael Brines, Patricio T. Huerta, Chunyan Li

## Abstract

**Aims:** This study aims to elucidate the underlying mechanisms of diving reflex, a powerful endogenous mechanism supporting underwater mammalian survival. Antioxidative responses, observed in marine mammals, may be contributing factors. Using a multi-organ approach, this study assesses whether acute and chronic diving reflex activate nuclear factor-erythroid-2-related factor 2 (NRF2) signaling pathways, which regulate cellular antioxidant responses.

**Methods:** Male Sprague-Dawley rats (*n*=38) underwent either a single diving session to elicit acute diving reflex, or daily diving sessions for 4-weeks to produce chronic diving reflex. NRF2 (total, nuclear, phosphorylated), NRF2-downstream genes, and malondialdehyde were assessed via Western blot, immunofluorescence, RT-PCR, and ELISA in brain, lung, kidney, and serum.

**Results:** Diving reflex increased nuclear NRF2, phosphorylated NRF2, and antioxidative gene expression, in an organ-specific and exposure time-specific manner. Comparing organs, the brain had the highest increase of phosphorylated NRF2 expression, while kidney had the highest degree of nuclear NRF2 expression. Comparing acute and chronic sessions, phosphorylated NRF2 increased the most with chronic diving reflex, but acute diving reflex had the highest antioxidative gene expression. Notably, calcitonin gene-related peptide appears to mediate diving reflex’ effects on NRF2 activation.

**Conclusions:** Acute and chronic diving reflex activate potent NRF2 signaling in the brain and peripheral organs. Interestingly, acute diving reflex induces higher expression of downstream antioxidative genes compared to chronic diving reflex. This result contradicts previous assumptions requiring chronic exposure to diving for induction of antioxidative effects and implies that the diving reflex has a strong translational potential during preconditioning and postconditioning therapies.

**Key Points:**

- Diving reflex activates potent NRF2 signaling via multiple mechanisms, including phosphorylation, nuclear translocation, and KEAP1 downregulation with both acute and chronic exposure.
- Diving reflex activates NRF2 via differential pathways in the brain and other organs; phosphorylated NRF2 increases more in the brain, while nuclear NRF2 increases more in the peripheral organs.
- Acute diving reflex exposure induces a more pronounced antioxidative effect than chronic diving reflex exposure, indicating that the antioxidative response activated by diving reflex is not dependent upon chronic adaptive responses and supports diving reflex as both a preconditioning and postconditioning treatment.

## 1. Introduction

The diving reflex (DR) is a powerful endogenous mechanism that enhances mammalian survival under hypoxic/anoxic conditions,^1–3^ partially by augmenting antioxidant defense responses, although the specific underlying mechanisms remain incompletely elucidated.^4–10^ DR is initiated by a combination of trigeminal sensory afferent activation and apnea, eliciting various physiological responses associated with antioxidative stress (**Fig. 1**).^1–3^ Notably, peripheral vasoconstriction and reflexive apnea, key physiological responses induced by DR, have demonstrated the capability to trigger the activation of nuclear factor-erythroid-2-related factor 2 (NRF2) signaling pathways, resulting in antioxidative stress effects.^11–17^ NRF2, a redox-sensitive transcription factor, plays a pivotal role in the antioxidant response system responsible for maintaining cellular homeostasis.^18,19^ The concurrent involvement of peripheral vasoconstriction and apnea within DR suggests the potential involvement of NRF2 in the antioxidative effects elicited by DR, although further investigation into this aspect is warranted. Given the historical successes of endogenous mechanisms in transitioning to therapeutic interventions,^20–22^ exploring the molecular mechanisms underlying DR may pave the way for its utilization as a non-invasive, non-pharmacological, multimodal therapy.

**Figure 1:**
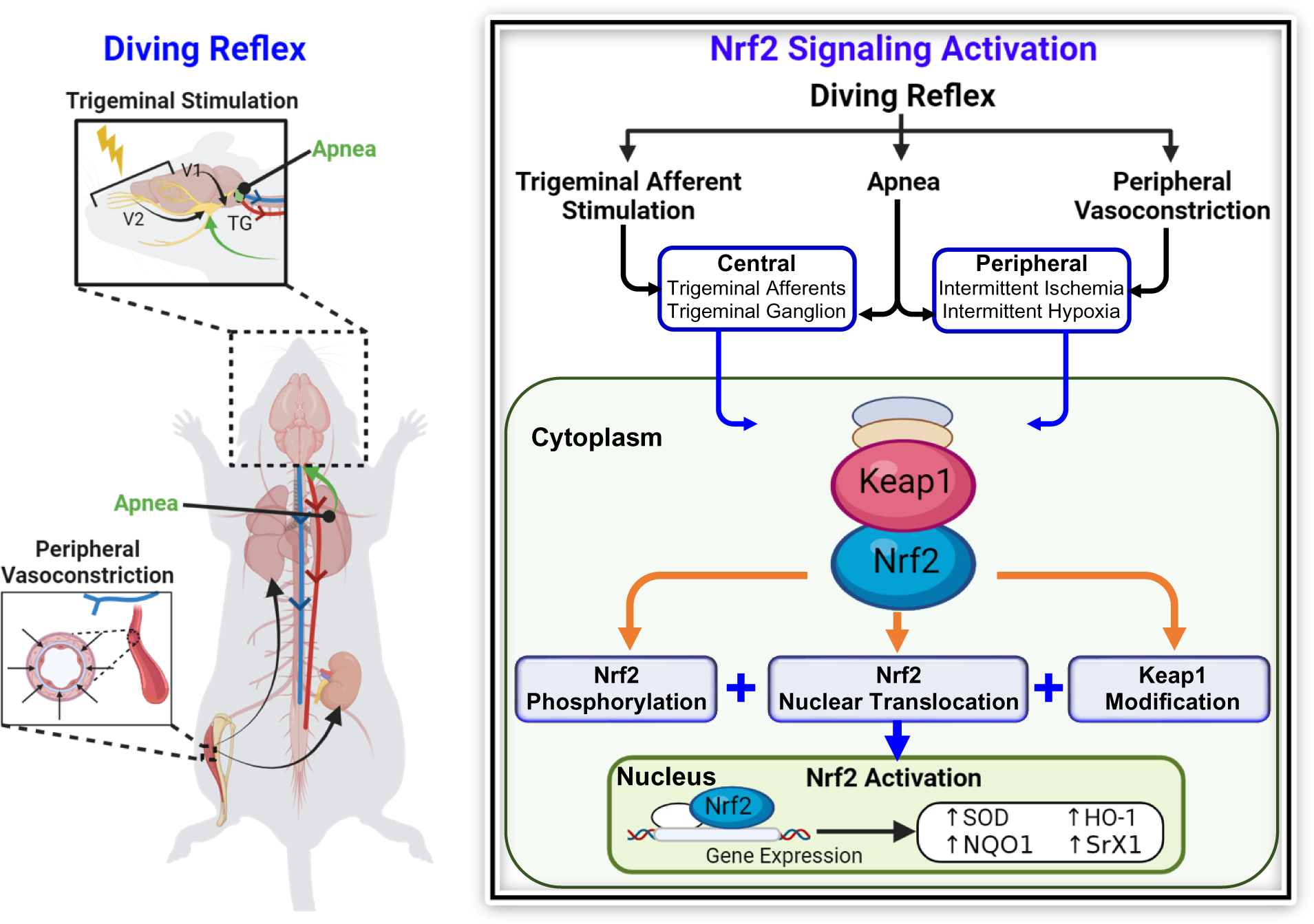
Factors underlying the actions of diving reflex on oxidative stress. Three major physiological effects contribute to the anti-oxidative effects of diving reflex: trigeminal stimulation, apnea, and peripheral vasoconstriction. Centrally, trigeminal afferent stimulation and apnea induce activation of the trigeminal afferents and ganglion, in turn inducing the release of neuropeptides. Apnea also induces upregulation of lactic acid in the peripheral organs, in conjunction with peripheral vasoconstriction. Neuropeptide release and peripheral lactic acid upregulation modulate the KEAP1–NRF2 complex, resulting in a combination of NRF2 phosphorylation, NRF2 nuclear translocation, and modification of KEAP1. Production of antioxidant enzymes and molecules is induced by activated NRF2. The different methods of NRF2 activation lends diving reflex a strong therapeutic potential for treatment of oxidative stress. Abbreviations: HO1, heme oxygenase 1; KEAP1, Kelch-like ECH-associated protein 1; NRF2, nuclear related factor 2; NQO1, NAD(P)H quinone dehydrogenase; SOD, superoxide dismutase; SRX1, sulfiredoxin 1 (Created with BioRender.com).

DR has demonstrated robust antioxidative effects in individuals engaged in chronic diving activities.^6–10^ Increased antioxidant enzyme activity and decreased reactive oxygen species (ROS) have been observed in chronic diving mammals.^8^ Frequent DR episodes in these animals correlate with lower levels of superoxide radical production. Similarly, trained human breath-hold divers show higher antioxidative enzyme activity levels following even a single diving session.^9^ Mechanistic explanations for these antioxidative effects assume a reliance on chronic adaptive responses resulting in increases of antioxidative enzyme activity levels.^6,7,9,10^ However, it is unknown whether chronic adaptation is required for DR to induce antioxidative effects and whether an acute diving session might have an antioxidative effect similar to repetitive diving. Furthermore, the potential link between the antioxidative effect observed in chronic diving^6–10^ and NRF2 signaling remains unexplored.

To assess whether DR activates NRF2-linked signaling pathways and produces antioxidative effects, we trained rats to dive through an underwater path using a well-established model of rodent DR (**Fig. 2**).^23–25^ Following training, rats underwent either a single session of intermittent diving, which we term ‘acute DR’ (ADR), or daily sessions of intermittent diving over the course of 4 weeks, which we term ‘chronic DR’ (CDR). Expression of NRF2 and other markers of oxidative stress were assessed in brain, kidney, lung, and serum (by Western blot, immunofluorescence, and RT-PCR methods) to assess possible mechanistic pathways. Our results indicate that following ADR and CDR, non-aquatic mammals experience a profound activation of the NRF2 signaling pathway connected to a potent increase of antioxidative genes. These findings imply that DR could hold therapeutic promise as a modulator of oxidative stress in both chronic and acute traumatic conditions.

**Figure 2:**
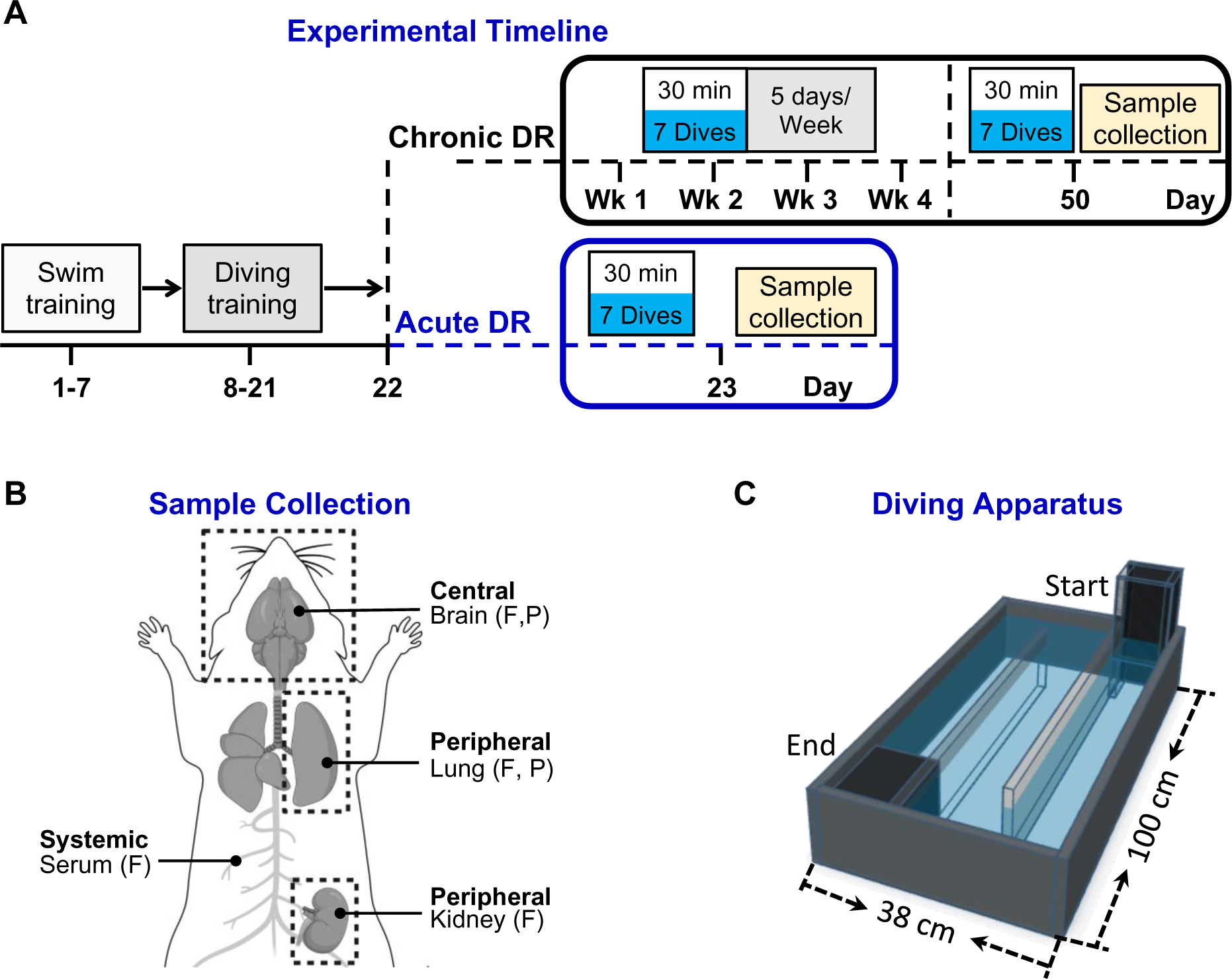
Conceptual diagram of the experimental protocol. **(A)** Timeline of dive training and diving reflex (DR) induction. Animals underwent 7 days of swim training, followed by 14 days of diving training. On Day 23, acute rats underwent a single 30-min session comprised of 7 dives, followed by 30-min of recovery and sample collection. Chronic rats underwent 4 weeks of daily diving 5 times per week, following the same 30-min dive session pattern as the acute rats. On day 50, the final diving session for the chronic rats was completed, again followed by 30-min recovery and sample collection. (**B**) Samples for both acute DR and chronic DR were collected from central and peripheral sources (F = fresh, P = perfused). **(C)** Schematic diagram of the diving tank. A modified version of the McCulloch diving tank was used which shortened the number of 1 m lanes from 5 to 3 and decreased the total distance from 5 m to 2.5 m. (Created with Biorender.com).

## 2. Results

### 2.1. DR modulates NRF2 nuclear translocation in the brain and peripheral organs

The protein levels of total NRF2 and nuclear NRF2 were measured in brain, kidney, and lung via Western blot (**Fig. 3**). In the brain, nuclear NRF2 significantly increased in the ADR and CDR groups, as compared to Sham (Sham: 1.0±0.3; ADR: 1.6±0.3, *P*<0.01; CDR: 1.5±0.3, *P*<0.05, one-way ANOVA followed by Tukey test, **Fig. 3A**), whereas total NRF2 did not differ between groups (Sham: 1.0±0.4; ADR: 1.1±0.2; CDR: 0.8±0.3, **Fig. 3A**). The kidney expressed a similar pattern of significantly higher nuclear NFR2 in the ADR and CDR groups (Sham: 1.0±0.3; ADR: 1.5±0.3, *P*<0.05; CDR: 1.5±0.4, *P*<0.05, **Fig. 3B**) but no difference of total NRF2 (Sham: 1.0±0.3; ADR: 1.3±0.5; CDR: 1.4±0.5, **Fig. 3B**). In the lung, nuclear NRF2 was only different in the CDR group (Sham: 1.0±0.3; ADR: 2.0±0.8; CDR: 2.1±0.8, *P*<0.05, **Fig. 3C**) whereas all groups were similar for total NRF2 (Sham: 1.0±0.4; ADR: 0.8±0.2; CDR: 1.1±0.4).

**Figure 3.**
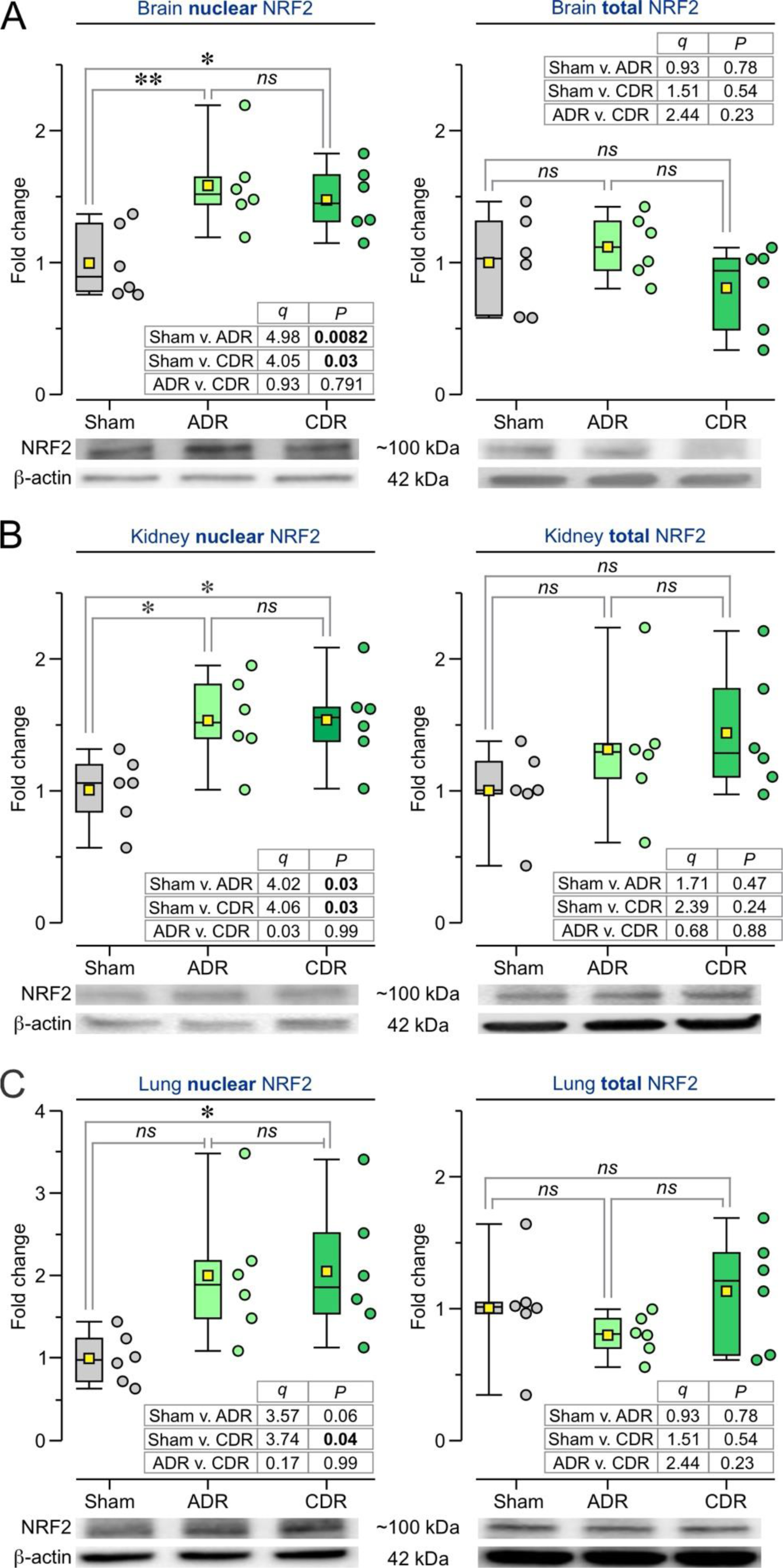
Diving reflex modulates NRF2 within the brain, kidney, and lung. NRF2 expression was measured within the brain, kidney, and lung using the Western blot method. Data are represented as boxplots (box = 25–75 range, line = median, yellow square = mean, whiskers = 10–90 range) and statistics from one-way ANOVA followed by Tukey test. (**A)** Brain samples show increased nuclear NRF2 expression for ADR (*N* = 6) and CDR (*N* = 6) rats compared to the Sham group (*N* = 6), and no statistical differences between groups for total NRF2 expression (right). **(B)** Kidney samples show a similar pattern as the brain, with higher nuclear NRF2 expression and unchanged total NRF2 in the ADR and CDR groups. **(C)** Lung samples also show elevated nuclear NRF2 expression and unaffected total NRF2 in the ADR and CDR groups. Abbreviations: *ns* = not significant, * *P* < 0.05, ** *P* < 0.01.

### 2.2. DR increases NRF2 phosphorylation in the brain and peripheral organs

Immunofluorescent staining indicated that DR increased the expression of phosphorylated NRF2 (pNRF2) in brain and lung (**Fig. 4**). We measured the intensity of the pNFR2 signals in several brain areas (**Fig. 4A**) and used two-way repeated ANOVA (with subject and brain region as repeated measures) followed by Tukey test for statistical analysis. pNRF2 increased significantly in the ADR and CDR groups within the dorsal cortex (Sham: 8.4±2.7; ADR: 15.2±3.3, *P*<0.001; CDR: 25.4±3.4, *P*<0.005), corpus callosum (Sham: 0.2±0.2; ADR: 5.1±1.3; CDR: 10.5±3.6, *P*<0.001), CA1 (Sham: 4.8±1.1; ADR: 17.2±2.3, *P*<0.001; CDR: 20.3±1.7, *P*<0.001), dentate gyrus (Sham: 10.9±6.5; ADR: 23.0±4.3, *P*<0.001; CDR: 34.7±6.4, *P*<0.001), amygdala (Sham: 3.8±1.0; ADR: 14.3±2.1, *P*<0.001; CDR: 21.4±5.2, *P*<0.001) and hypothalamus (Sham: 11.6±2.5; ADR: 25.2±4.5, *P*<0.001; CDR: 30.3±6.6, *P*<0.001). Within the lung, pNRF2 increased 3-to-6-fold in the ADR and CDR groups, in relation to Sham (Sham: 3.8±2.0; ADR: 9.6±3.2, *P*<0.001; CDR: 17.7±2.8, *P*<0.001, one-way ANOVA with Tukey test, **Fig. 4B**). Post-hoc comparison between the ADR and CDR groups revealed that CDR produced significantly higher pNRF2 expression in brain and lung as compared to ADR (**Fig. 4**), although there was no difference in nuclear NRF2 levels.

**Figure 4.**
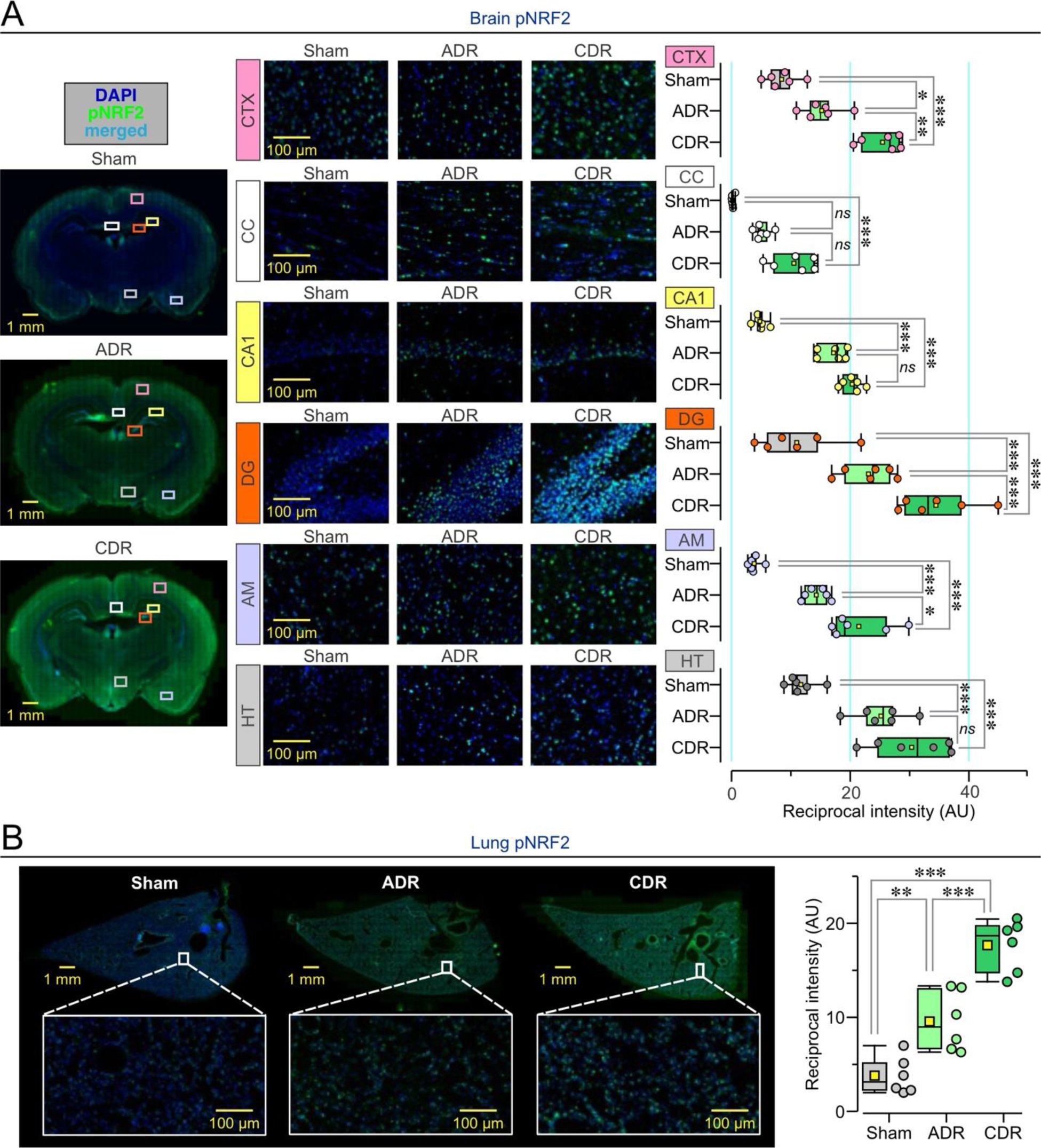
Diving reflex increases phosphorylated NRF2 expression. **(A)** *Left,* pNRF2 expression (green color) within the brain was assessed using immunofluorescent staining. DAPI (blue color) was used to stain cell nuclei. The dorsal cortex (CTX), corpus callosum (CC), CA1 region of the hippocampus (CA1), dentate gyrus (DG), amygdala (AM), and hypothalamus (HT) illustrate increased pNRF2 in the ADR (*N* = 6) and CDR (*N* = 6) brains compared to Sham brains (*N* = 6). *Right,* data are represented as boxplots (box = 25–75 range, line = median, yellow square = mean, whiskers = 10–90 range). Significantly increased pNRF2 expression is observed in the ADR and CDR brains. (**B**) pNRF2 expression within the lung increases with both ADR and CDR. Abbreviations: *ns* = not significant, * *P* < 0.05, ** *P* < 0.01, *** *P* < 0.001 (repeated-measures ANOVA with Tukey test).

### 2.3. DR promotes modification of KEAP1 in the brain and peripheral organs

The mRNA levels of genes associated with NRF2 activation pathways, including Kelch-like ECH associated protein 1 (KEAP1) and p62, were measured in brain and lung (**Fig. 5**). p62 interrupts the NRF2–KEAP1 protein-protein interactions to activate NRF2 without oxidation.^19^ Within the brain, KEAP1 expression decreased significantly in both DR groups compared to Sham (Sham: 1.0±0.1; ADR: 0.8±0.1, *P*<0.01; CDR: 0.7±0.1, *P*<0.005, one-way ANOVA with Tukey test, **Fig. 5A**), whereas p62 expression showed significant increase in DR groups (Sham: 0.9±0.1; ADR: 1.1±0.1, *P*<0.005; CDR: 1.1±0.05, *P*<0.05, **Fig. 5B**). In the lung, KEAP1 expression decreased significantly in the ADR group only (Sham: 1.1±0.1; ADR: 0.8±0.2, *P*<0.05; CDR: 0.9±0.05, **Fig. 5C**), while p62 expression did not show significant change (Sham: 1.0±0.2; ADR: 0.8±0.1; CDR: 1.0±0.07, **Fig. 5D**).

**Figure 5:**
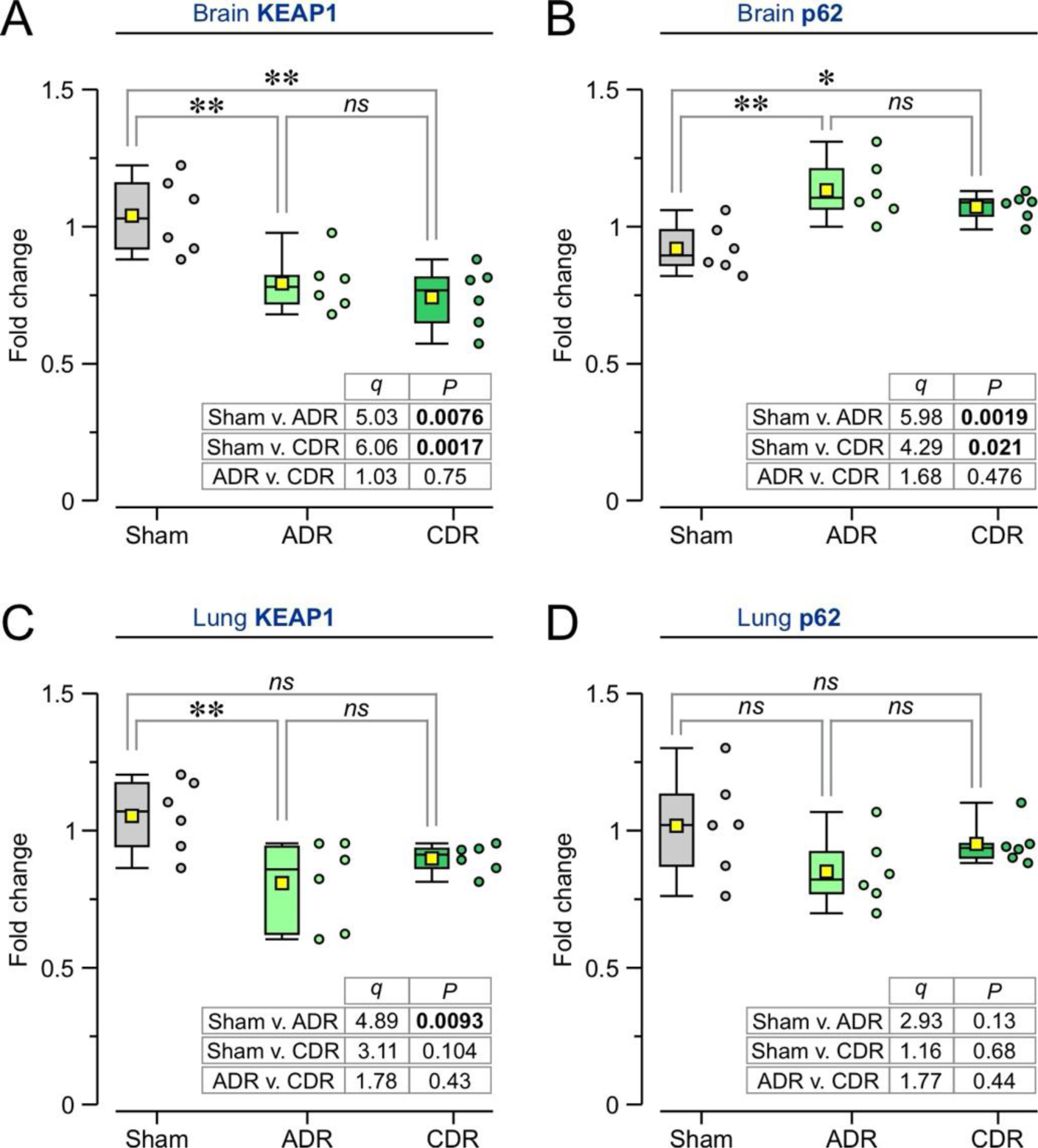
Diving reflex modulates differential NRF2 activating factors. KEAP1 RNA and p62 RNA were measured within the brain and lung. Data are represented as boxplots (box = 25–75 range, line = median, yellow square = mean, whiskers = 10–90 range) and statistics from one-way ANOVA followed by Tukey test. (**A)** Brain samples show decreased RNA expression for KEAP1 in ADR (*N* = 6) and CDR (*N* = 6) rats compared to the Sham group (*N* = 6). **(B)** Brain p62 is elevated in the ADR and CDR groups. **(C)** Lung KEAP1 expression is decreased in the ADR group only. **(D)** Lung p62 expression is similar across all groups. Abbreviations: *ns* = not significant, * *P* < 0.05, ** *P* < 0.01.

### 2.4. DR increases antioxidant enzyme activity without increasing oxidative stress

The mRNA levels of NRF2 downstream genes, including heme-oxygenase-1 (HO1), superoxide dismutase (SOD), NAD(P)H-quinone-dehydrogenase-1 (NQO1), and sulfiredoxin-1 (SRX1) were assessed in brain and lung (**Fig. 6**). For the brain (**Fig. 6A**), ADR and CDR groups showed significant increases of HO1 (Sham: 1.0±0.1; ADR: 2.0±0.3, *P*<0.001; CDR: 1.5±0.2, *P*<0.01, one-way ANOVA with Tukey test), SOD (Sham: 1.0±0.1; ADR: 1.3±0.1, *P*<0.001; CDR: 1.2±0.04, *P*<0.005), NQO1 (Sham: 1.0±0.1; ADR: 1.6±0.3, *P*<0.001; CDR: 1.4±0.1, *P*<0.01) and SRX1 (Sham: 1.1±0.2; ADR: 1.8±0.3, *P*<0.001; CDR: 1.3±0.2). Within the lung (**Fig. 6B**), ADR and CDR groups had significant increases of HO1 (Sham: 1.0±0.1; ADR: 2.0±0.3, *P*<0.001; CDR: 1.5±0.3, *P*<0.05), SOD (Sham: 1.0±0.1; ADR: 1.9±0.2, *P*<0.001; CDR: 1.5±0.2, *P*<0.001), NQO1 (Sham: 1.1±0.1; ADR: 1.3±0.1, *P*<0.005; CDR: 1.4±0.1, *P*<0.001), and SRX1 (Sham: 1.0±0.2; ADR: 2.2±0.6, *P*<0.001; CDR: 1.6±0.4, *P*<0.05). Interestingly, the post-hoc tests showed that the ADR group had significantly higher expression levels of HO1, SOD, NQO1 and SRX1 than the CDR group (**Fig. 6**).

**Figure 6:**
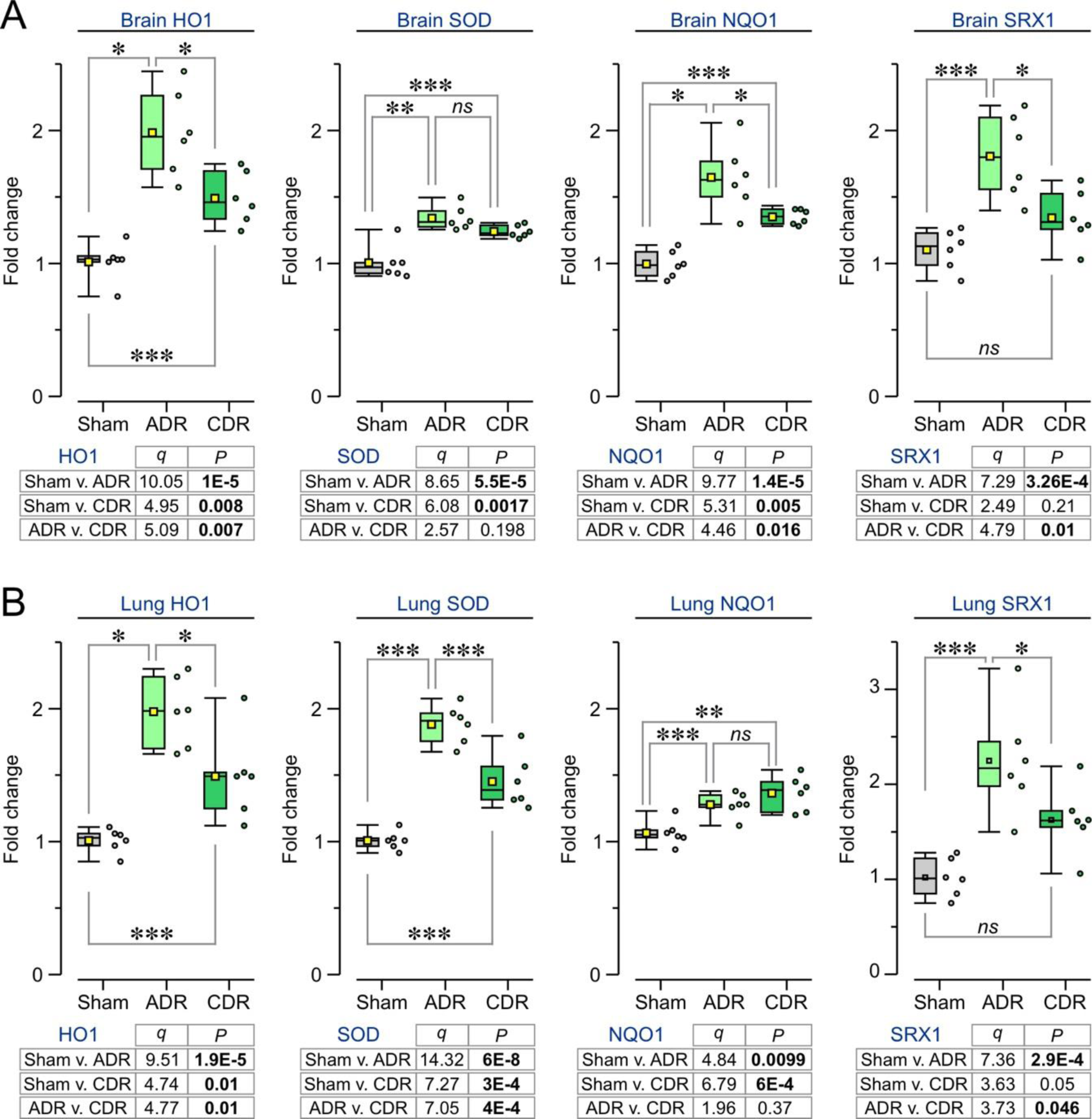
Diving reflex increases antioxidative gene expression. Assessment of the mRNA levels of several NRF2 downstream genes, such as heme-oxygenase-1 (HO1), superoxide dismutase (SOD), NAD(P)H-quinone-dehydrogenase-1 (NQO1), and sulfiredoxin-1 (SRX1) in brain and lung. Data are represented as boxplots (box = 25–75 range, line = median, yellow square = mean, whiskers = 10–90 range) and statistics from one-way ANOVA followed by Tukey test. (**A)** Brain samples show significant elevations of HO1, SOD, NQO1 RNA expression in the ADR and CDR groups; SRX1 RNA is higher in the ADR group only. (B) The lung also shows significant increases of HO1, SOD, NQO1 RNA expression in the ADR and CDR groups; SRX1 RNA is higher in the CDR group only. Abbreviations: *ns* = not significant, * *P* < 0.05, ** *P* < 0.01, *** *P* < 0.001.

We also assessed lipid peroxidation via measurement of malondialdehyde in the serum, brain, lung, and kidney via colorimetric means. Malondialdehyde concentration decreased in the CDR brain, CDR kidney and ADR kidney. The changes were not statistically significant in the serum (Sham: 1.0±0.3, ADR: 0.9±0.4, CDR: 1.1±0.3), brain (Sham: 0.05±0.02, ADR: 0.05±0.01, CDR: 0.04±0.01), lung (Sham: 0.03±0.01, ADR: 0.03±0.01, CDR: 0.03±0.01), and kidney (Sham: 0.06±0.01, ADR: 0.05±0.01, CDR: 0.05±0.01).

### 2.5. DR activates CGRP in the brain and other organs

Previous assessments have shown that calcitonin gene-related peptide (CGRP) can increase the expression of NRF2.^26,27^ As such, we measured the CGRP levels by Western blot in brain, lung, and kidney after DR exposure and found that CGRP was higher in the ADR and CDR groups (**Fig. 7**) for the brain (Sham: 1.0±0.4; ADR: 3.7±1.9, *P*<0.005; CDR: 3.0±1.4, **Fig. 7A**), kidney (Sham: 1.0±0.6; ADR: 1.6±0.3, *P*<0.05; CDR: 1.8±0.3, *P*<0.005, **Fig. 7B**), and lung (Sham: 1.0±0.3; ADR: 2.0±0.8, *P*<0.05; CDR: 2.4±0.8, **Fig. 7C**). Interestingly, the brain expression of receptor activity modulating protein-1 (RAMP1), a CGRP receptor subunit, was significantly increased in the CDR but not the ADR group (Sham: 1.0±0.2; ADR: 1.4±0.3; CDR: 1.8±0.3, *P*<0.0001, **Fig. 7D**). This defines an important distinction between acute and chronic diving exposure.

**Figure 7:**
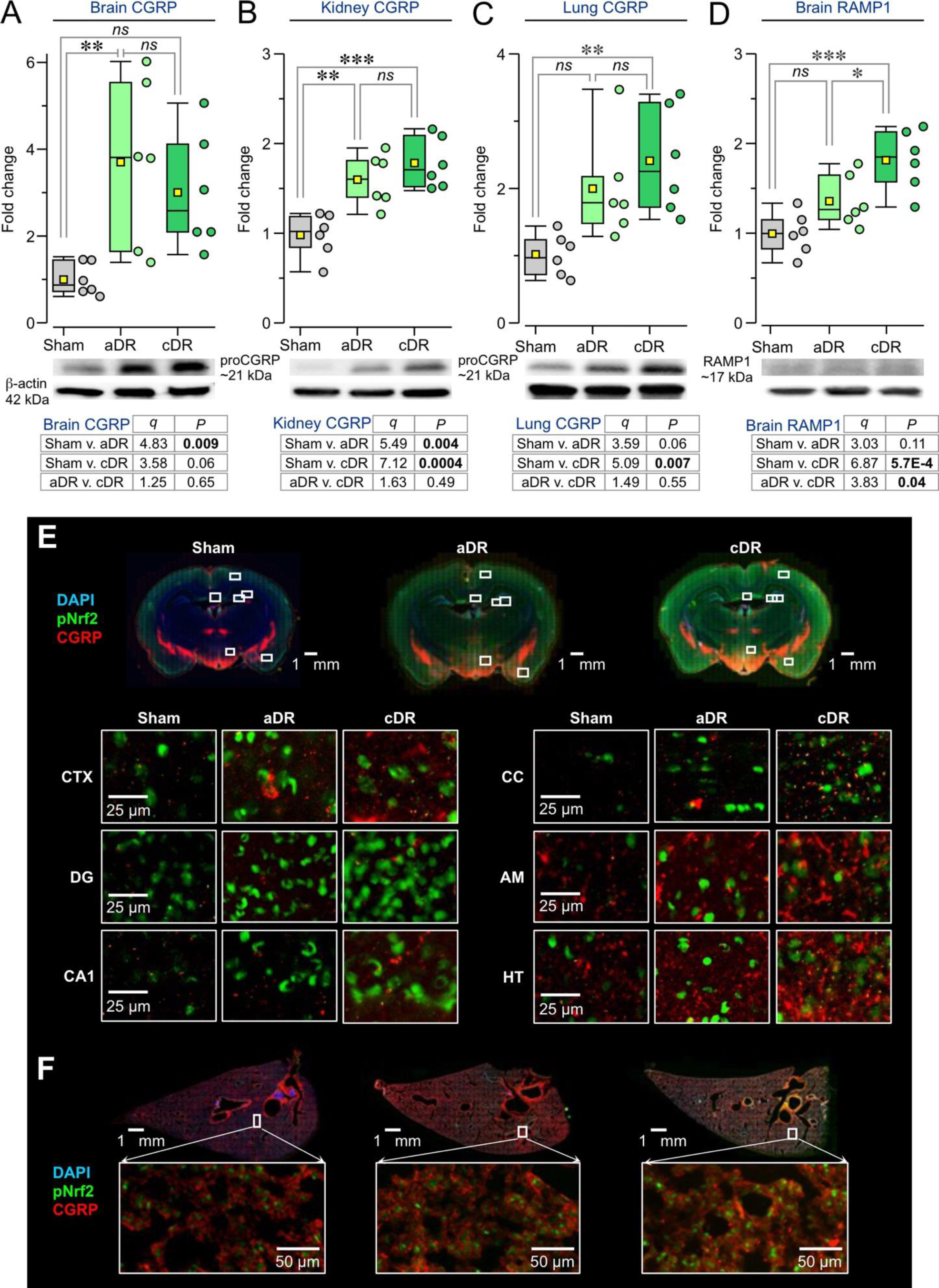
Diving reflex promotes NRF2 phosphorylation through CGRP within the brain, kidney, and lung. **(A-D)** CGRP and RAMP1 expressions were assessed within the brain, kidney, and lung using the Western blot method. Data are represented as boxplots (box = 25–75 range, line = median, yellow square = mean, whiskers = 10–90 range) and statistics from one-way ANOVA followed by Tukey test. **(A)** Brain CGRP expression is increased in the ADR group only. **(B)** Kidney CGRP expression is increased in the ADR and CDR groups. **(C)** Lung CGRP expression is increased in the CDR group only. **(D)** Brain RAMP1 expression is also increased exclusively in the CDR group. **(E)** CGRP (red color) and pNRF2 expressions (green color) were assessed by immunofluorescent staining in the brain. Visually, for the ADR and CDR groups, increased expression of both intracellular and extracellular CGRP correlates with more intense pNRF2 expression in the dorsal cortex (CTX), corpus callosum (CC), CA1 field of the hippocampus (CA1), dentate gyrus (DG), amygdala (AM), and hypothalamus (HT). **(F)** Immunofluorescent staining in the lung also shows correlated staining of CGRP and pNRF2. Abbreviations: *ns* = not significant, * *P* < 0.05, ** *P* < 0.01, *** *P* < 0.001.

Immunofluorescent staining of region-specific CGRP expression showed enhanced colocalization between CGRP and pNRF2 expression (**Fig. 7E, F**). This included the dorsal cortex (Sham: 6.0±2.1; ADR: 18.7±6.2; CDR: 25.6±3.2), corpus callosum (Sham: 1.3±0.8; ADR: 3.2±0.9; CDR: 10.8±4.5), CA1 (Sham: 1.6±0.4; ADR: 3.2±1.3; CDR: 13.2±5.1), dentate gyrus (Sham: 5.9±5.1; ADR: 16.9±4.1; CDR: 19.8±4.1), amygdala (Sham: 11.9±5.5; ADR: 25.4±4.1; CDR: 28.1±9.7), and hypothalamus (Sham: 11.8±2.8, ADR: 24.8±5.0; CDR: 29.6±7.1) as well as the lung (Sham: 13.6±7.6; ADR: 34.9±5.8; CDR: 41.5±3.0) (**Supplementary Figure 1**).

## 3. Discussion

Our findings illustrate that DR triggers a robust activation of NRF2 signaling in a time- and organ-specific manner. Both acute and chronic diving exposures elicit NRF2 signaling activation through a combination of NRF2 phosphorylation, NRF2 nuclear translocation, and KEAP1 downregulation. Remarkably, the ADR group exhibits increased expression of antioxidative genes relative to their CDR counterparts, concomitant with the absence of further escalation in nuclear NRF2 levels. These observations imply alterations in redox sensitivity and a shift in NRF2 signaling between acute and chronic exposures. Furthermore, peripheral organs exhibit a greater enhancement of nuclear NRF2 translocation, while the brain displays a more pronounced increase in NRF2 phosphorylation, indicating divergent pathways of NRF2 activation between the brain and peripheral organs. In conjunction with the intrinsic ability of DR to trigger NRF2 signaling without exacerbating oxidative stress, our study underscores that both acute and chronic exposures induce robust activation of NRF2 signaling pathways, leading to potent antioxidative responses, thereby positioning DR as a promising translational therapeutic intervention for pre- and post-conditioning.

A key aspect of our results is the time-specific dependance of DR exposure. While nuclear NRF2 expression does not differ between the ADR (∼1.3-2X) and CDR (∼1.3-2.8X) groups, phosphorylated NRF2 expression in the brain and lung is significantly higher in CDR rats (∼2.5-9X) compared to ADR rats (∼1.5-7X). This suggests that prolonged exposure to DR may predispose NRF2 signaling to phosphorylation. However, NRF2 needs to translocate to the nucleus to regulate the transcription of antioxidant genes.^28^ *In vitro* investigations have demonstrated that while phosphorylated NRF2 tends to translocate to the nucleus, a fraction remains in the cytoplasm.^29^ NRF2 is a redox-sensitive gene transcription factor and contains multiple motifs regulating its distribution between the nucleus and cytoplasm. Under oxidative stress conditions, the motifs responsible for nuclear export become inactive, favoring NRF2 translocation.^30^ However, under conditions of heightened resistance to oxidative stress (such as CDR), the sensitivity of nuclear exporting motifs diminishes, resulting in a disparity between the rate of production and translocation of phosphorylated NRF2.^30^ Consequently, chronic exposure likely induces epigenetic alterations that diminish redox sensitivity and modulate NRF2 activation, necessitating further investigation.

We observed more pronounced expression on antioxidative genes during acute diving than chronic diving. Specifically, ADR induced an ∼1.3-2.2X upregulation of antioxidative genes within the brain and lung, while CDR resulted in an ∼1.2-1.6X upregulation. Interestingly, this coincided with a non-significant decrease in malondialdehyde levels in the ADR kidney and CDR brain and kidney (∼0.2X), suggesting the maintenance of antioxidative gene activity. In contrast, both dimethyl fumarate (DMF) and sulforaphane, as NRF2 activators, exhibit little to no effect on lipid peroxidation in the lung,^31^ liver,^32–34^ and smooth muscle^35^ of healthy rodents, but induce a non-significant yet noticeable increase in malondialdehyde levels in the kidneys.^31,36^ This contradicts the effects observed with DR. Experimentally, hypoxia-reoxygenation cycles generate ROS and reactive nitrogen species, inducing cellular cytolytic injury even without inflammatory cells present.^37^ Diving mammals, however, display lower levels of superoxide radical production despite repeated exposure to hypoxia and reoxygenation cycles.^8^ This is regardless of the fact that DR itself generates an initial hypoxia.^2,3^ Animals engaging in more frequent dives exhibit elevated levels of antioxidant enzymes, such as SOD and glutathione reductase, indicating a dose-dependent effect of DR on resistance to hypoxic stress. Additionally, professional breath-hold divers experience decreased post-exercise blood lactic acid build-up, acidosis, and lipid peroxidation, indicating reduced oxidative stress overall.^9^ Chronic divers also demonstrate an ∼16% increase in SOD activity levels relative to non-divers when exposed to DR.^7^ These findings collectively suggest an augmentation of antioxidant enzyme activity and an adaptive resilience to antioxidant depletion induced by DR,^6–9^ consistent with the observed antioxidative gene expression and malondialdehyde levels in this experiment. This underscores the potent increase in antioxidative gene expression induced by DR, which is accompanied by an exposure time-dependent epigenetic enhancement of antioxidant enzyme activity.

The epigenetic modulation of antioxidant enzyme activity corresponds with the impact of DR on CGRP activity. Both ADR and CDR significantly elevated CGRP levels in the brain, lung, and kidney. In the brain, the principal contributor is trigeminal sensory afferent activation, which directly releases stored CGRP from trigeminal sensory afferent nerve endings into the surrounding tissue.^38–40^ Conversely, the increase in CGRP levels in the kidney and lung may stem from peripheral nerve exocytosis of CGRP mediated by intermittent hypoxia- and ischemia-induced acidosis.^41–45^ While trigeminal activation leads to systemic CGRP elevation, as evidenced by an ∼2-3X increases in jugular CGRP levels during cluster headache attacks,^46^ the extent to which it affects peripheral organs remains uncertain. Systemically, CGRP is endogenously released as a protective response to increased oxidative stress, associated with NRF2 activation.^26.27.46–58^ It has been observed to boost SOD production by ∼80% and increase NRF2 and HO1 expression by 80% and 70%, respectively, concomitant with a return of ROS production and malondialdehyde levels to baseline in an *in vitro* high glucose model.^49^ CGRP also augments SOD levels in Schwann cells,^46^ aligning with the observed increase in SOD by ADR. Moreover, in a mouse model of traumatic brain injury, CGRP has been shown to upregulate p62 via protein kinase C mediation,^59,60^ consistent with the observed increase in p62 and NRF2 activation herein.^61^ The principal limiting factor for CGRP, however, is the RAMP1 receptor subunit.^62,63^ The approximately 1.9X increase in RAMP1 expression observed with CDR, but not ADR, suggests that although both groups exhibit comparable CGRP expression levels, CDR animals likely have higher overall CGRP activity. Through its influence on protein kinase A, adrenocorticotropin,^64^ and progesterone,^65,66^ the sustained upregulation of CGRP, as observed within CDR, establishes a positive feedback loop leading to increased RAMP1 expression. This, in turn, amplifies overall CGRP activity, correlating with the elevation in NRF2 phosphorylation observed with CDR. Consequently, CDR enhances overall CGRP activity and NRF2 activation, likely contributing to a shift in the predisposition of the NRF2 signaling pathway. In summary, the initial upregulation of CGRP by DR modulates the initial antioxidative effect of DR, while the sustained effect of DR on RAMP1 and CGRP activity influences the epigenetic modulation of NRF2 phosphorylation and antioxidant enzyme activity.

The findings presented here also illustrate that DR activates robust Nrf2 signaling pathways in a manner specific to individual organs. DR potently activates NRF2 signaling across the brain, resulting in ∼1.4X increase in nuclear NRF2 expression and ∼1.5X-7X increase in phosphorylated NRF2 expression. These changes are comparable to the ∼2X increase in total NRF2 observed in the aged rat hippocampus following treatment with DMF, an FDA-approved pharmacologic NRF2 activator.^36,67,68^ Moreover, one of the physiological effects of DR, intermittent hypoxia alone, induces a substantial increase in total NRF2 (∼3X) in an *in vitro* blood-brain barrier model.^11^ This finding aligns with the observed increase in phosphorylated NRF2 in the hippocampus by ADR (2–4X). However, the rise in hypoxia-induced phosphorylated NRF2 is coupled with a simultaneous elevation in ROS levels, approximately threefold higher.^11^ Furthermore, DR increases both nuclear and phosphorylated NRF2 levels in peripheral organs (∼2-3X), comparable to the increase in total NRF2 levels in the kidney induced by DMF (∼2.8X). Nevertheless, a comparison between the effects of DR on the brain and peripheral organs reveals that the brain experiences a greater increase in Nrf2 phosphorylation (∼1.5-7X vs. ∼3X), whereas peripheral organs experience a greater increase in nuclear Nrf2 translocation (∼1.4X vs. ∼2X). This suggests organ-specific activation of Nrf2 signaling pathways by DR. Overall, our experimental results demonstrate that DR induces a potent activation of Nrf2 signaling pathways in multiple organs, rendering it an appealing option for therapeutic intervention.

Findings from both the brain and peripheral organs suggest that DR enhances NRF2 signaling pathways through various mechanisms involving modifications of KEAP1. DR induces an ∼0.2-0.3X reduction in KEAP1 levels in both the brain and lung, suggesting the occurrence of KEAP1 bond oxidation.^69^ KEAP1-mediated activation of NRF2 signaling involves various mechanisms.^19,28^ Oxidation of KEAP1 bonds releases NRF2, promoting its translocation to the nucleus. However, the pharmacological strategy targeting KEAP1 oxidation lacks specificity and may inadvertently oxidize additional molecules, resulting in unintended outcomes. For instance, DMF functions by oxidizing KEAP1 bonds, thereby promoting NRF2 activation and reducing KEAP1 expression.^69^ DR, however, also induces an ∼1.2X increase in p62 levels in the brain. p62, along with molecules such as p21, Wilms tumor gene, and dipeptidyl peptidase III, exhibits a preferential binding affinity for KEAP1 over NRF2.^19,28^ These molecules directly disrupt protein-protein interactions involving KEAP1, leading to enhanced release and aggregation of NRF2 upon their upregulation. This mechanism demonstrates fewer off-target effects compared to oxidative activation of the NRF2 signaling pathway. The concurrent decrease in KEAP1 levels in the brain and lung, along with the elevation of p62 in the brain, suggests that DR may enhance NRF2 activation through both oxidative and non-oxidative means. This dual mechanism of NRF2 activation sets DR apart from oxidative NRF2 activators like DMF^19,28,69^ and may contribute to a synergistic effect in NRF2 signaling pathway activation, potentially increasing its versatility.

This study has several limitations. Firstly, the assessment of antioxidative efficacy occurred only 30-min following DR. Although the findings herein suggest a robust antioxidative effect associated with both acute and chronic DR exposures, the sustainability of this effect, particularly in the case of acute DR, remains uncertain. In chronically diving humans it was possible to measure a decrease in oxidative stress markers within the blood 15-h following final DR induction, indicating a durable antioxidative response,^10^ and suggesting avenues for further examination. The assessment of NRF2 activation at multiple post-DR timepoints will be valuable for applying DR to acute injuries in which oxidative stress plays a major role. Secondly, we only investigated CGRP as one of the factors facilitating NRF2 phosphorylation. However, it is widely recognized that trigeminal activation releases various other antioxidative neuropeptides in addition to CGRP.^40,70^ These neuropeptides could potentially influence NRF2 signaling and serve as underlying factors contributing to the effects of DR.^40,70^ Thirdly, the pathways affecting oxidative stress are complex. In this study, we only focused on the NRF2 pathway. Accordingly, additional investigation is needed to unravel the mechanisms underlying the impact of DR on oxidative stress.

## 4. Materials and Methods

### 4.1. Experimental animals

This study was approved by the Institutional Animal Care and Use Committee of the Feinstein Institutes for Medical Research and performed in accordance with the National Institutes of Health *Guidelines For The Use Of Experimental Animals*. Male Sprague-Dawley rats weighing 100–125g at the beginning of the experiment were used, and all rats were between 350–450g at the time of collection (Charles River Laboratories, New York, USA). Animals were housed in a temperature-controlled room (12-h light/dark cycle), in cages lined with Enrich-o’Cobbs bedding (The Anderson, Inc, Maumee, Ohio) and were given access to food and water *ad libitum*.

### 4.2. Experimental groups

A total of 38 animals were obtained as a single cohort (Charles River Laboratories, New York, USA), of which 26 underwent training into adulthood (**Fig. 2**). Animals were excluded based off their performance and behavior during training for result consistency (N=2). Following training, successful animals were divided into the ADR (*N*=12) and CDR (*N*=12) groups. The remaining animals (*N*=12) aged up as comparative Sham.

### 4.3. Rat voluntary diving protocol

Rats were trained to voluntarily dive through a multi-channel underwater tunnel, for a total length of 2.5 m, using a modified version of the method laid out by McCulloch et al.^23^ and confirmed by McCulloch^24^ and Hult et al.^25^ Rats were required to show swimming ease within the first two days of training, voluntarily initiate dives within the first week of diving training, and display minimal signs of agitation when handled or diving. Following successful completion of training, ADR rats underwent a final 30-min diving session, comprised of repeated dives every 5-min, for a total of 7 dives, followed by sample collection (**Fig. 2**). Rats in the CDR group underwent 30-min DR trials 5 times per week for a period of 4-weeks. On Day 50, they underwent a final 30-min diving session and sample collection. Additional training details are described in **Supplementary Materials 1**.

### 4.4. Sample collection and analysis

Samples from animals in the ADR, CDR, and sham groups were obtained by a combination of either fresh collection or supradiaphragmatic transcardial perfusion (N=6 per group). Western blot and immunofluorescence staining methods were used to assess total NRF2, nuclear NRF2, phosphorylated NRF2, CGRP, and RAMP1 expression. NRF2 activation and downstream effectors were assessed by RT-PCR for HO1, SOD, NQO1, Srx1, KEAP1, and p62. Lipid peroxidation was assessed via ELISA for malondialdehyde. Detailed experimental procedures are described in **Supplementary Materials 1**.

### 4.5. Statistical analysis

Western blot and rtPCR data are presented as fold change respective to sham and immunofluorescence data are represented as reciprocal intensity (AU). All data are presented as mean±SD and represented visually as box-and-whisker plots, with mean and individual data points present. A power calculation was used (α=0.05 and β=0.8), which indicated a minimum group size of *N*=6. The normal distribution of each variable was verified using both the Shapiro–Wilk test and the Kolmogorov–Smirnov tests. Significant differences between the three groups were assessed using a one-way analysis of the variance test (ANOVA) followed by Tukey post-hoc test. The immunofluorescent data (used in **Fig. 4**) was analyzed with two-way repeated ANOVA (with subject and brain region as the repeated measures) followed by Tukey test. Data were considered significantly different with *α*<5% (*p*<0.05). All statistical analyses were performed using GraphPad Prism (version 9, GraphPad Software, Boston, MA) and OriginPro (version 2022b 64-bit SR1, OriginLab Corp., Northampton, MA).

## 5. Conclusions

The findings presented in this study demonstrate that both acute and chronic DR activation robustly stimulate the NRF2 signaling pathway, leading to a pronounced antioxidative effect in both brain and peripheral organs. DR induces NRF2 phosphorylation, facilitates NRF2 nuclear translocation, and downregulates KEAP1 expression, resulting in distinct time- and organ-specific activation patterns. The temporal specificity of NRF2 signaling activation is associated with an adaptive adjustment of redox sensitivity and antioxidant gene expression upon chronic DR exposure. Notably, the activation of NRF2 by DR operates via a dual mechanism of KEAP1 modification, conferring DR with additional therapeutic potential compared to pharmacological NRF2 activators that act through singular mechanism. These findings represent an initial step towards elucidating the precise mechanisms underlying DR’s modulation of oxidative stress, laying the groundwork for future clinical therapeutic applications.

## Author Contributions

**K. Powell**: Investigation; formal analysis; methodology; visualization; writing – original draft preparation; writing – review & editing.

**S. Wadolowski**: Investigation; methodology; writing – review & editing.

**W. Tambo:** Investigation; methodology; writing – review & editing.

**J.J. Strohl:** Formal analysis; writing – review & editing.

**D. Kim:** Methodology; writing – review & editing.

**J. Turpin**: Writing – review & editing.

**Y. Al-Abed**: Writing – review & editing.

**M. Brines**: Writing – review & editing.

**P.T. Huerta**: Formal analysis; methodology; visualization; writing – review & editing.

**C. Li**: Conceptualization; investigation; formal analysis; methodology; visualization; writing – original draft preparation; writing – review & editing; funding acquisition; project administration; resources; supervision.

## Acknowledgements

### Data Availability Statement

Individual data points are plotted for all graphs. The data that support the findings of this study are available from the corresponding author upon reasonable request.

## Ethics Approval Statement

Experiments within this study were approved by the Institutional Animal Care and Use Committee of the Feinstein Institutes for Medical Research and performed in accordance with the National Institutes of Health *Guidelines For The Use Of Experimental Animals*.

