## Supplemental Materials 1 for "Intrinsic diving reflex induces potent antioxidative response by activation of NRF2 signaling"

### Supplemental Materials and Methods

**Voluntary Diving Training:** Rats were trained to voluntarily dive and swim through a multi-channel underwater tunnel using a modified version of the method laid out by McCulloch et al. [23] and confirmed by McCulloch[24] and Hult et al.[25] Rats were trained within a rectangular tank (100 x 38 x 15 cm) comprised of ¾-in thick acrylic pieces, with a ½-in thick top (MazeEngineers, Skokie, IL). Over the course of 1-week the rats were acclimated to the water environment and trained to voluntarily swim a 3-channel route for a total distance of ~2.5m. This training consisted of first developing the rats' familiarity and comfort with the water environment and then gradually increasing the distance which they are expected to traverse, with a total of 2.5m at a depth of about 10cm in 32°C water. Rats were trained daily, with 3-5 trials per day per rat. Each rat was placed in the water at increasing distances from the finish platform. When the rats showed consistent successful negotiation of the path, they started on dive training to last another 2-weeks. It consisted of first developing the rats' ability to dive underwater comfortably and voluntarily, and then gradually increasing the length of the underwater portion of the swimming path, until they showed consistent negotiation of the underwater multi-channel tunnel.

**Fresh Sample Collection and Processing:** At the time of final sample collection, animals were heavily anesthetized using isoflurane and blood samples were collected via the vena cava (N=6/group). Blood samples were refrigerated (4°C) for 30 min, and then spun down for 10 min at 2000G in order to collect the resulting serum. The same animals were decapitated and samples were collected from the brain, kidney, and lung.  Frozen tissue samples were powdered in liquid nitrogen and stored until usage at -80°C.

**Transcardial Perfusion and Processing:** A separate set of animals (N=6/group) was heavily anesthetized and supradiaphragmatically transcardially perfused using 0.1M ice-cold phosphate buffered saline followed by 4% paraformaldehyde. The head was removed by guillotine and the brains and lungs were collected. Samples were placed in 4% paraformaldehyde overnight, followed by graded sucrose solution, and cryopreserved in a mix of 30% sucrose and optimal cutting temperature media (Electron Microscopy Sciences, USA), and stored at -80°C until sectioning. Brains and lungs were coronally cryosectioned at a thickness of 14 um on a cryostat (Leica Biosystems, Germany), mounted on Polysine glass slides (Thermo Fisher Scientific, Waltham, MA), and stored at -80 °C until staining.

**Western Blot:** Powdered tissue samples were aliquoted from brain, lung, and kidney. For assessments of nuclear Nrf2 (nNrf2), a commercial kit was used for isolation of nuclear protein (Abcam, Waltham, MA). Assessment of total protein utilized tissue homogenized in RIPA lysis buffer (Thermo Fisher Scientific, Waltham, MA) containing protease and phosphatase inhibitor cocktail (Thermo Fisher Scientific, Waltham, MA) with a bead mill homogenizer. In both cases, protein concentration was quantified using the BCA protein assay kit (Thermo Fisher Scientific, Waltham, MA). Protein samples were separated on a 4-20% TGX gel (Bio-Rad, Hercules, California) by electrophoresis, according to molecular weight, using the Protein Plus Kaleidoscope (Bio-Rad, Hercules, California) for visualization. Proteins were then electro-transferred onto polyvinylidene difluoride membranes using the semi-dry transfer method, blocked with either 5% skimmed milk (Thermo Fisher Scientific, Waltham, MA) (CGRP (Santa Cruz Biotechnology, Dallas, Texas), RAMP1 (Proteintech, Rosemont, IL)) or 5% bovine serum albumin (Millipore Sigma, Burlington, MA) (Nrf2 (Proteintech, Rosemont, IL)) in 1X Tris-buffered saline with Tween

20 (TBST) (Millipore Sigma, Burlington, MA) at room temperature for 1 hour and then incubated with primary antibodies overnight at 4°C. After three washes in TBST, the membranes were incubated with respective HRP-conjugated secondary antibodies (Abcam, Waltham, MA) in 5% milk at room temperature for 1 hour, followed by another three washes with TBST. Signals were detected by chemiluminescence using ECL substrate (Thermo Fisher Scientific, Waltham, MA) on a Bio-Rad ChemiDoc Imaging System. ImageJ was then used to quantify relative protein levels in the blots. Any changes to brightness or contrast for the purpose of visualization were consistent across entire blots. β-actin (Sigma, Burlington, MA) and Lamin B1 (Proteintech, Rosemont, IL) were used as loading controls for total and nuclear protein, respectively. All Western Blot data is expressed as a ratio of sham.

**Malondialdehyde Measurement:** Lipid peroxidation was measured within brain, kidney, and lung powdered tissue samples, as well as serum samples, via assessment of malondialdehyde (Lipid Peroxidation (MDA) Assay Kit (Colorimetric/Fluorometric), Abcam, Waltham, MA). Tissue samples were processed according to high protein protocols.

**RT-PCR:** Total RNA from powdered brain and lung tissue was isolated using Trizol reagent (Life Technologies, Carlsbad, CA). High-capacity cDNA Reverse Transcription Kit (Applied Biosystems, Foster City, CA) was used to synthesize cDNA from the isolated RNA. RT-PCR was performed using the primer sequences as follows: (1) heme oxygenase-1 (F:ACAGGGTGACAGAAGAGGCTAA; R:CTGTGAGGGACTCTGGTCTTTG), (2) superoxide dismutase (F:GCTCTAATCACGACCCACT; R:CATTCTCCCAGTTGATTACATTC) , (3) NQO1 (F:GCGTCTGGAGACTGTCTGGG; R:CGGCTGGAATGGACTTGC), (4) Srx1 (F:CCCAAGGCGGTGACTACTAC; R:GGCAGGAATGGTCTCTCTCTGTG), (5) Keap1 (F:GGACGGCAACACTGATTC; R:TCGTCTCGATCTGGCTCATA), and (6) p62 (F:TCCCTGTCAAGCAGTATC C; R:TCCTCCTTGGCTTTGTCTC). All primers were obtained from Eurofins Genomics (Louisville, KY). qPCR was performed on a 7500 Real-time PCR System (Applied Biosysems, Foster City, CA) utilizing SYBR Green PCR Master Mix reagents (Applied Biosystems, Foster City, CA). Rat GAPDH (F:AGG TTG TCT CCT GTG ACT TC; R:CTG TTG CTG TAG CCA TAT TC) was used as an endogenous control. The delta-delta calculation method was utilized to obtain fold change relative to controls.

**Immunofluorescence Staining:** Cryosectioned Polysine were washed with TBST, blocked with 5% goat serum (Abcam, Waltham, MA) supplemented with 1% bovine serum albumin (Sigma, Burlington, MA) for 1 hour at room temperature and sequentially incubated with primary antibody, mouse anti-CGRP antibody (Santa Cruz Biotechnology, Dallas, Texas) at a dilution factor of 1:50, at 4°C overnight and its corresponding secondary antibody, goat anti-mouse HRP (1:400), at room temperature for 1 hour. The slides were then co-stained with rabbit anti-phosphorylated Nrf2 antibody (Abcam, Waltham, MA) at a dilution factor of 1:100, and its corresponding secondary antibody, goat anti-rabbit HRP (1:400), at room temperature for 1 hour. Slides were counterstained with DAPI (1:2000, Thermo Fisher Scientific, Waltham, MA) and mounted with Vectashield Antifade mounting medium (Vector Laboratories, Burlingame, CA). The EVOS M7000 imaging system (Thermo Fisher Scientific, Waltham, MA) was used to visualize and image the slides, using the 20× objective and automated XY-stitching function to obtain whole brain images. For the quantification of signal expression semi automatically, single-channel images were assessed in ImageJ.[71,72] The range of pixel intensities of images was between 0 and 255, representing the weakest and strongest intensity, respectively.

**SUPPLEMENTAL FIGURE 1**

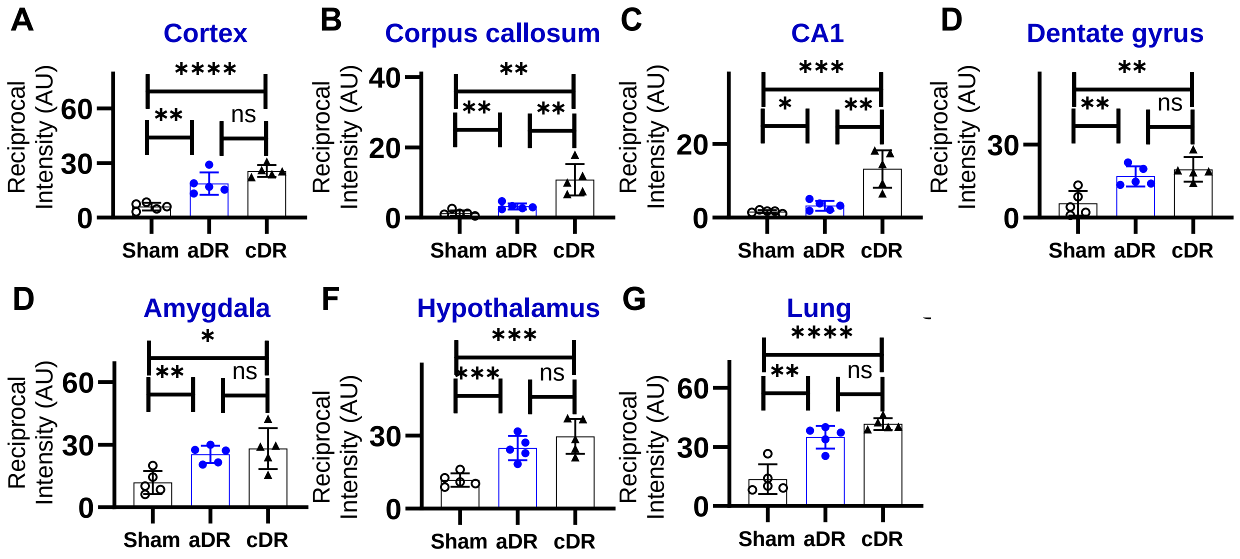


**Supplemental Figure 1. Diving reflex increases CGRP expression.** CGRP and pNRF2 expressions were assessed by immunofluorescent staining in the midbrain and lung. Regions of increased CGRP expression correlate with regions of increased pNrf2 expression. (**A-G**) CGRP expression intensity increases with both aDR and cDR. Abbreviations: ns=not significant, *=p<0.05, **=p<0.01, ***=p<0.001, ****=p<0.0001.
